# AniMarkerDB: a comprehensive database for exploring cell types and marker genes in livestock and poultry at single-cell resolution

**DOI:** 10.1101/2025.10.14.682327

**Authors:** Zhuohang Li, Tao Zhang, Xueqing Li, Jiangwu Huang, Zimin Xie, Fei Gao, Haiming Cai, Mingfei Sun, Manman Dai, Ming Liao

## Abstract

Single-cell RNA sequencing (scRNA-seq) has dramatically advanced the understanding of cellular heterogeneity. While numerous marker gene databases are available for humans and mice, a lack of systematic resources for livestock and poultry species remains, limiting progress in functional genomics, immunology, and breeding.. To address this challenge, we developed AniMarkerDB (https://animarkerdb.bio), a comprehensive and curated database dedicated to marker genes and immune-related epitopes in economically animals, including chicken, pig, and duck. AniMarkerDB integrates 7,010 marker gene across 37 tissues and 846 cell types, together with 71,442 immune epitope records from IEDB. All entries undergo rigorous literature curation, manual validation, and multi-level quality control, with standardized nomenclature and annotation to ensure data consistency and reusability. The platform supports flexible queries by species, tissue, cell type, or gene. It offers analytical tools for cross-species comparison model organisms such as human and mouse, interactive single-cell atlas visualization, and user-defined cell type annotation. Additionally, AniMarkerDB provides dynamic visualizations and export options, enabling researchers to efficiently obtain large-scale marker and epitope data for downstream applications such as infectious disease research, vaccine target design, and comparative immunology. Looking ahead, AniMarkerDB will expand species coverage and incorporate additional modalities, including single-cell atlases from healthy and disease models, establishing itself as a comprehensive and authoritative platform for animal cell biology, disease modeling, and translational research.

**Graphical abstract:** 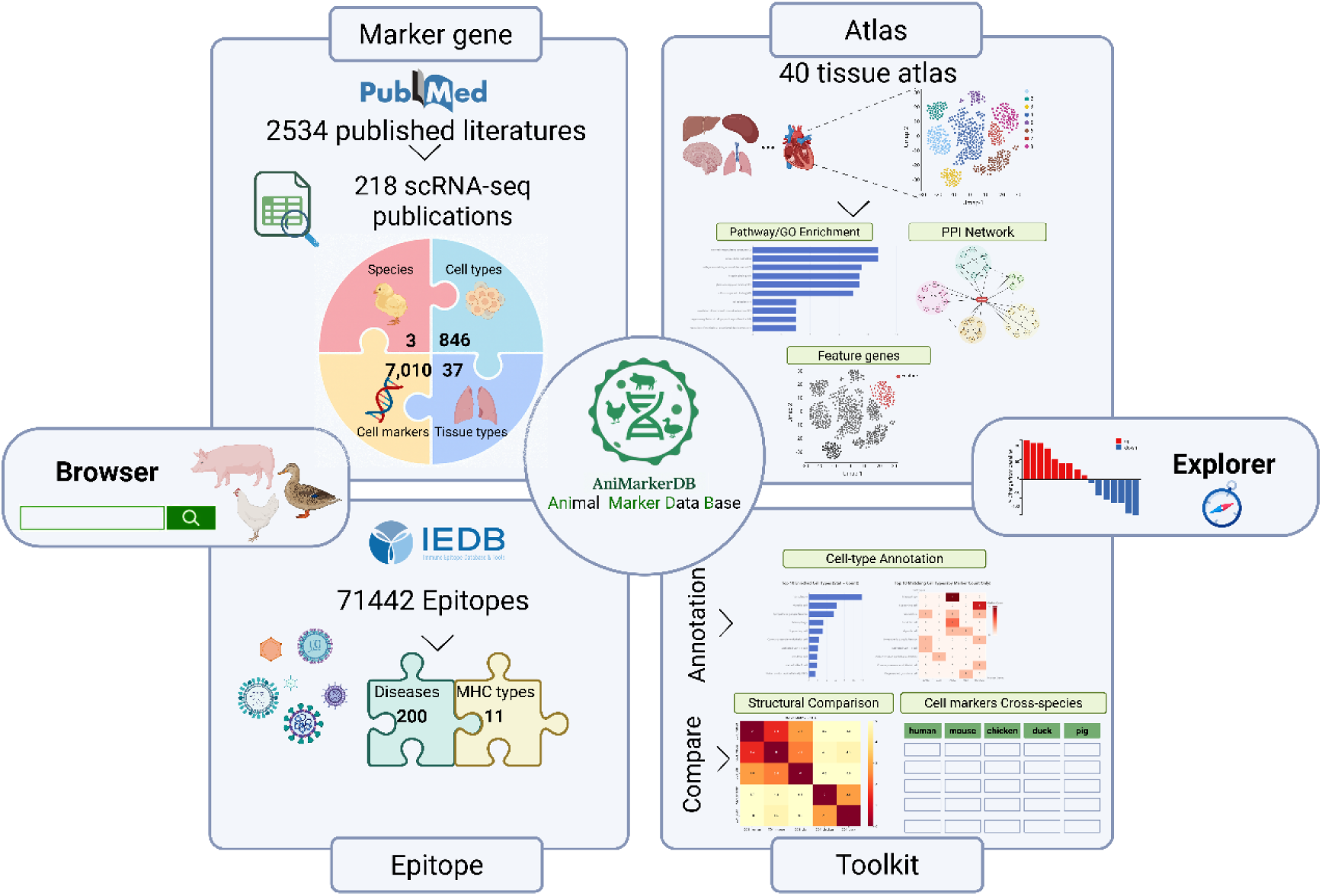

## INTRODUCTION

Single-cell transcriptomics has transformed biological research by enabling high-resolution profiling of cellular diversity across various tissues, species, and developmental stages(1,2). While the earliest applications were primarily confined to human and model organisms, scRNA-seq has been increasingly adopted in livestock and poultry research (3). In particular, studies in species such as chicken, pig, and duck have yielded unprecedented insights into cellular heterogeneity, tissue architecture, and immune system function(3–5). Mohammadinejad *et al*. employed transcriptomic profiling to identify regulatory genes and pathways crucial for skeletal muscle growth in cattle, sheep, and pigs, thereby highlighting genetic markers that are valuable for livestock breeding and production improvement (6). These economically essential animals not only serve as vital sources of food and agricultural products but have also emerged as valuable models for studying genetics, immunology, and disease mechanisms. For instance, single-cell immune profiling in chickens and ducks has identified novel immune cell types and deepened our understanding of avian resistance mechanisms(7). In the context of xenotransplantation, Saad-Bay *et al*. employed single-cell RNA and TCR sequencing to elucidate the dynamics of donor-reactive T cells and innate immune cells during pig-to-human organ transplantation, identifying new targets for immunosuppressive therapy (8). Additionally, spatial transcriptomic analysis has been applied to study liver pathology following pig-to-primate islet transplantation, uncovering pathways related to fatty liver disease and adipogenesis (9). Recent advances demonstrate how emerging sequencing technologies and computational approaches now enable researchers to systematically dissect gene expression at single-cell resolution across a wide range of tissues and cell types (4). Together, these advances have laid the groundwork for precise cell type definition, lineage tracing, and in-depth exploration of immune mechanisms in livestock species.

Despite these advances, significant challenges remain in the field of single-cell research for livestock and poultry. First, information on marker genes and cell type annotation is highly dispersed across the literature, often buried in supplementary files with inconsistent data formats and nomenclature standards(10,11). This fragmentation hinders cross-tissue and cross-species comparisons, reproducible cell-type annotation, and broader functional analysis. While databases such as CellMarker(12), CellMarker 2.0(13), PanglaoDB(14), and scPlantDB(15) have provided structured resources for human, mouse, and plant cell markers, available resources for livestock and poultry are still limited in terms of species and tissue coverage, annotation consistency, and functional depth. Furthermore, the limited availability of commercial antibodies and the incomplete functional annotation of many marker genes in these species pose additional challenges for experimental validation and immune studies(16–18).

To address these limitations, we developed AniMarkerDB (https://animarkerdb.bio), a manually curated and standardized database of marker genes and immune epitopes for chicken, pig, and duck. The database integrates high-quality data from 132 single-cell studies, along with 71,442 epitope records retrieved from the IEDB. AniMarkerDB offers flexible query capabilities, cross-species marker exploration, structure-based similarity prediction, pathway enrichment tools, and interactive single-cell atlas visualization. This platform aims to provide a centralized, authoritative resource to accelerate research in livestock and poultry cell biology, immunity, and disease modeling.

## MATERIAL AND METHODS

### Data collection and literature curation

To systematically build AniMarkerDB — a specialized marker gene database for chicken (*Gallus gallus*), pig (*Sus scrofa*), and duck (*Anas platyrhynchos*) — we curated single-cell transcriptomic studies published since 2019 from the PubMed database. Search strategies incorporated keywords such as “single-cell RNA sequencing,” “cell type identification,” and “marker gene discovery”. Studies were screened based on explicit cell type definitions and reported marker genes. Review articles, studies lacking cell annotation information, or those with incomplete metadata were excluded. Accession numbers from eligible studies (*e.g*., GEO, SRA, CRA) were collected to facilitate subsequent data integration.

From each publication, we extracted the species, tissue origin, annotated cell types, marker genes, sequencing platform information, accession IDs, and bibliographic details. All marker genes were standardized using the NCBI Entrez E-utilities to retrieve official gene symbols, Entrez Gene IDs, protein IDs, and functional descriptions(19). Synonymous gene names were unified following official gene symbols. As of July 31, 2025, AniMarkerDB includes 98 chicken studies, 113 pig studies, and 7 duck studies, encompassing 7,010 marker records across 37 tissues and 846 cell types.

### Data processing and matrix standardization

To construct comprehensive single-cell transcriptomic atlases for livestock species, we applied a unified data processing and matrix standardization pipeline to all curated datasets. In total, AniMarkerDB integrates 2,224,866 chicken cells and 3,861,087 pig cells, originating from 17 chicken tissues and 24 pig tissues, and covering major biological systems such as the immune, digestive, reproductive, and muscular systems. Datasets were primarily obtained from public repositories, including GEO (Gene Expression Omnibus), SRA (Sequence Read Archive), CRA (Genome Sequence Archive), CNGBdb (China National GeneBank DataBase), and USDA (United States Department of Agriculture).

For datasets providing ready-to-use sparse matrices (*e.g*..mtx or .h5 formats), we directly incorporated them into downstream analyses without additional preprocessing. For projects offering only raw sequencing data (*e.g*., SRA format or FASTQ files), a standardized preprocessing pipeline was applied. Raw reads were aligned to the appropriate reference genomes (*GRCg7b* for chicken and *Sscrofa11.1* for pig), followed by feature-barcode matrix generation. Different sequencing platforms were processed using dedicated pipelines: CellRanger (v9.0.0) was applied for 10x Genomics datasets, while Celescope2 (v2.4.0) was used for BGI GEXSCOPE and DNBSEQ C4/C5 datasets. All resulting expression matrices were standardized into .h5ad format to facilitate uniform downstream integration and analysis. Datasets originating from the same tissue type and species were subsequently integrated using Scanpy (v1.10.3)(20). To ensure consistent gene annotation across datasets, gene identifiers were unified based on Ensembl IDs and corresponding official gene symbols(21).

### Quality control and batch effect correction

All single-cell datasets were preprocessed using a standardized Scanpy (v1.10.3)(20) pipeline to ensure data quality and cross-sample consistency. Cells with <200 detected genes or mitochondrial gene expression >10% were excluded to remove low-quality cells and technical artifacts. To further eliminate potential doublets, Scrublet (v0.2.3)(22) was applied to compute doublet scores, and cells with scores >0.3 were filtered out.

Following quality control, batch effect correction was performed using the scVI model (scvi-tools, v1.2.0) (23), which models cellular transcriptomic features in a latent space to harmonize expression distributions while preserving biological heterogeneity. The effectiveness of batch correction was evaluated by calculating Silhouette coefficients (24) based on latent embeddings using scikit-learn (v1.5.2) (23), thereby quantifying the extent of batch mixing. Subsequently, principal component analysis (PCA) was conducted on the latent representations to extract the top 30 principal components, followed by the construction of a k-nearest neighbor (kNN) graph for mapping cell relationships. Cell clusters were then identified using the Leiden(24) community detection algorithm, starting with an initial resolution of 1.0 and dynamically adjusting the resolution to ensure that the number of clusters fell within the range of 10 to 50 and that each cluster contained at least 50 cells. Finally, Uniform Manifold Approximation and Projection (UMAP) was employed for the two-dimensional visualization of the global distribution of cell types (25).

### Cell-type annotation and functional enrichment analysis

Following clustering, differential expression analysis was performed for each cell cluster using the Wilcoxon rank-sum test, as implemented in Scanpy (v1.10.3) (20). Filtering criteria included a gene detection frequency >25% within the cluster, log2 fold change >1, and an adjusted P-value <0.001 using the Benjamini-Hochberg method. The top 200 upregulated genes per cluster were selected to construct cluster-specific expression profiles. Cell type annotation was conducted by comparing these cluster-specific profiles against reference profiles curated in AniMarkerDB. Only reference expression profiles with an average expression level greater than 1 were considered for annotation to ensure reliability. Assignments were made based on the highest correlation scores, following a strategy inspired by expression profile-matching approaches used in SingleR (28), scmap (29), and CellTypist (26), ensuring robust cross-dataset annotation consistency.

Functional enrichment analysis was performed on signature genes for each identified cell type. Gene Ontology (GO) enrichment across biological process (BP), molecular function (MF), and cellular component (CC) categories, along with Kyoto Encyclopedia of Genes and Genomes (KEGG) pathway analysis, were conducted using g: Profiler (27) and clusterProfiler (v4.0.0)(28), with an FDR threshold of <0.05. Gene set enrichment analysis (GSEA) was performed using clusterProfiler (v4.0.0)(28), with reference gene sets obtained from MSigDB (v7.2)(29) to assess activation trends. Protein–protein interaction (PPI) networks were constructed using STRING(30) with a minimum confidence score threshold of 0.4.

### Epitope data integration

To supplement immune-related resources, experimentally validated epitope data for livestock and poultry species (primarily chicken, pig, and duck) were collected from the Immune Epitope Database (IEDB)(31). The records included antigen information, epitope amino acid sequences, associated MHC molecule classes, and origin. All epitope data were standardized and integrated into the database’s retrieval module, allowing users to filter results based on species, pathogen, or epitope sequence. To enhance the structural context, external links to corresponding MHC molecule entries were incorporated from the IPD-MHC database, enabling users to explore the spatial conformations of MHC-epitope binding interfaces (32).

### User-defined cell-type annotation

To enhance interactivity and flexibility, we implemented a user-defined annotation module that accepts custom marker gene sets uploaded by users. Cell type predictions are generated based on an internally curated marker database using two complementary strategies. First, a hypergeometric enrichment analysis evaluates the significance between user-submitted genes and reference marker sets, with composite scores calculated by weighting P-values and the number of matched genes. The top 10 predicted cell types are ranked and presented in a bar plot. Second, direct overlaps between user-provided markers and reference cell types are quantified to construct a matching matrix, which is visualized as a heatmap for intuitive interpretation. To ensure accurate matching, user-uploaded genes must conform to standardized gene symbol formats, and internal normalization is applied to correct case sensitivity and formatting inconsistencies.

### Search and cross-species comparison

AniMarkerDB provides an integrated search system to facilitate the efficient retrieval of cell marker information. Users can search by cell type name or gene symbol through a unified search interface on the homepage. Search results are displayed in sortable and filterable tables, including species, tissue, cell type, marker gene, and supporting references. The platform also promotes reverse lookup by marker gene, enabling users to query associated cell types based on specific gene inputs. Search outputs are structured with tabular presentations and intuitive visualizations, including summary tables of matched markers and word cloud figures highlighting the most representative marker genes based on the number of supporting occurrences.

Building upon the basic search functionality, AniMarkerDB incorporates a cross-species marker gene comparison module to enhance the interpretability and comparative utility of cell type annotation. Upon querying any cell type from chicken, pig, or duck, the platform simultaneously displays the Top 5 most frequently reported marker genes for the corresponding cell type across five species: chicken, pig, duck, human, and mouse. Marker gene information for humans and mice is sourced from CellMarker 2.0 (13), with cross-species matching based on standardized cell type nomenclature and unified gene symbol annotation. This module enables users to rapidly assess conserved marker expression patterns across species, providing valuable references for translational research, evolutionary studies, and functional validation. For marker genes with the same name across multiple species, AniMarkerDB allows users to assess structural similarities and differences across species. Protein structure data are retrieved from AlphaFoldDB(33), and root mean square deviation (RMSD) values are computed to quantify structural divergence. This module enables users to rapidly assess conserved marker expression and structural patterns across species, providing valuable references for translational research, evolutionary studies, and functional validation.

### Web interface and statistical visualization

AniMarkerDB utilizes its integrated search system, allowing for intuitive data visualization and exploration. The homepage provides an overview of species, tissues, and cell types covered in the database. For organ-level single-cell atlases, the platform presents tissue-specific UMAP projections, enabling users to explore the distribution of cell types within each tissue. Hovering over a specific cell cluster reveals real-time information including cell type annotation, source publication, and sample attributes. Additionally, a statistical module provides dynamic summaries of database contents, such as sample counts, cell type distributions, and marker gene frequencies, with interactive filtering by species and tissue origin. A detailed user guide is available to assist users in navigating search, visualization, and enrichment analysis workflows.

### Data availability

AniMarkerDB is freely available through the official website (URL to be provided upon release), offering unrestricted access to all integrated marker gene data and organ-level single-cell atlases. Users can browse, search, and download standardized datasets without the need for registration. All resources have been curated and organized in accordance with the FAIR principles (Findable, Accessible, Interoperable, and Reusable) to support data reuse and secondary development (34).

## Results

### Overview of AniMarkerDB

AniMarkerDB is the first comprehensive database dedicated to integrating multi-dimensional information on marker genes and immune epitopes in poultry and livestock. Focusing on three major economic animals—chicken, pig, and duck—AniMarkerDB systematically curates and integrates high-quality data from primary literature indexed in major databases such as PubMed, establishing a standardized data management and annotation framework. As of July 2025, AniMarkerDB contains 218 original publications, covering over 37 tissue types and 846 cell types, with thousands of marker gene entries (Figure 1), including 2,555, 4,250, and 205 entries for chicken, pig, and duck, respectively. All data are rigorously manually curated, consistently named, and subjected to multi-level quality control, with standardized gene symbols, protein IDs, and comprehensive metadata to ensure high accuracy and reusability across datasets and species.

**Figure 1.**
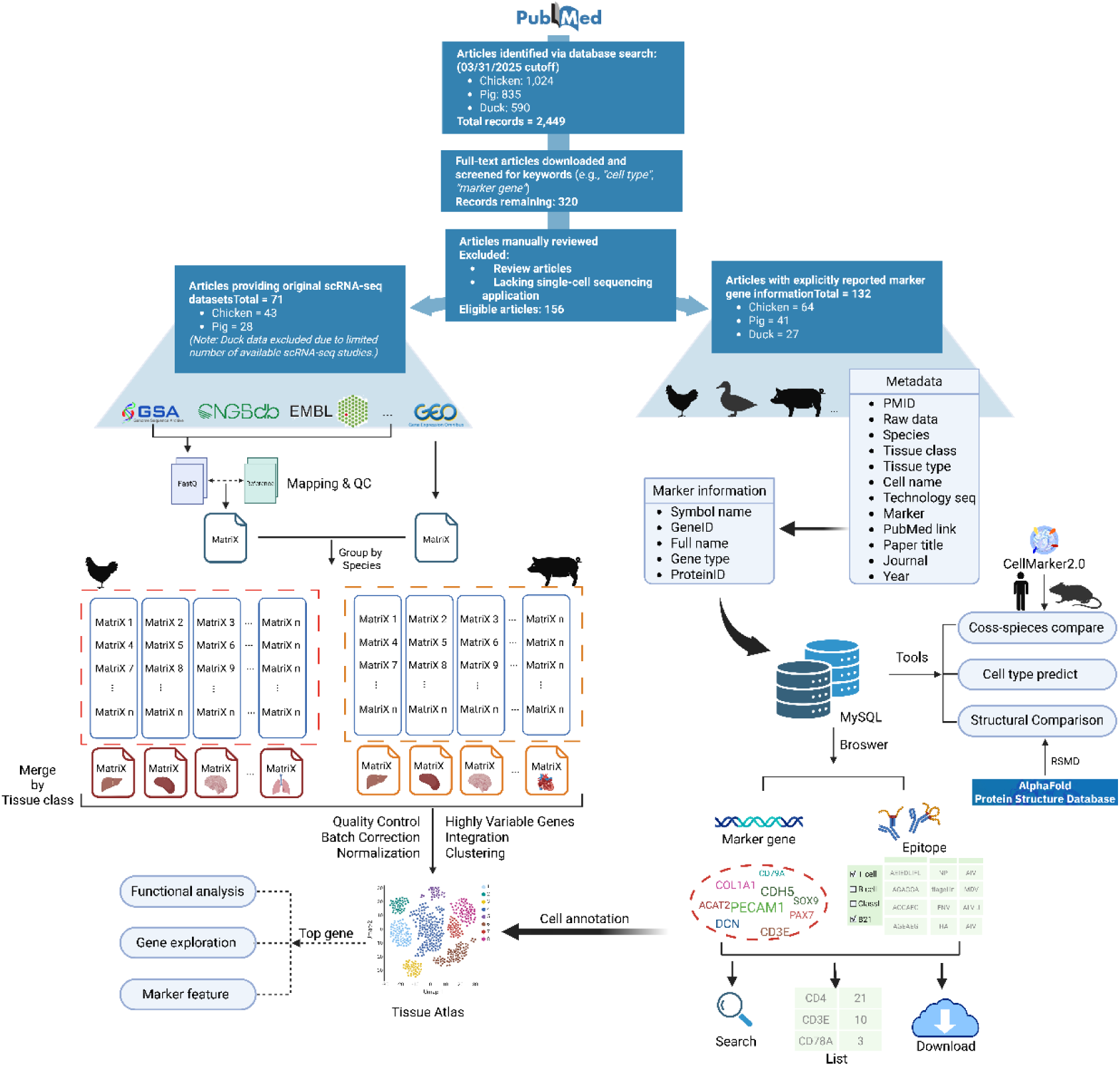
The architecture of AniMarkerDB.

The annotation framework of AniMarkerDB is compatible with mainstream resources, such as CellMarker 2.0 (13). Each marker gene entry includes detailed metadata, such as the gene’s full name, Gene ID, Protein ID, PMID, tissue, cell type, and sequencing platform (Figure 1), providing a solid foundation for downstream comparative and functional analyses. In parallel, the database systematically integrates immune epitope data from IEDB and compiles statistics on epitope distribution across species, cell types, and pathogens (31).

A core feature of AniMarkerDB is the standardized integration of primary scRNA-seq datasets from repositories such as GEO, GSA, EMBL, and others, enabling unified normalization and clustering across diverse tissue types. Automated cell-type annotation and hierarchical clustering facilitate the construction of comprehensive single-cell atlases for various tissues and cell populations. In total, we collected 2,224,866 chicken cells and 3,861,087 pig cells (Table 1), with most tissues contributing tens to hundreds of thousands of cells. Batch-correction performance was quantified using Silhouette Scores across major tissues; values were close to zero for most tissues, indicating effective removal of batch effects and consistent intra-tissue clustering. This extensive cell coverage, combined with robust quality control, establishes a solid foundation for constructing downstream single-cell atlases and identifying marker genes. Building upon these atlases, AniMarkerDB integrates functional enrichment analyses including GSEA, GO, KEGG, and PPI network reconstruction, thereby enabling in-depth investigation of cellular heterogeneity and biological function across species and tissue contexts.

**Table 1.**
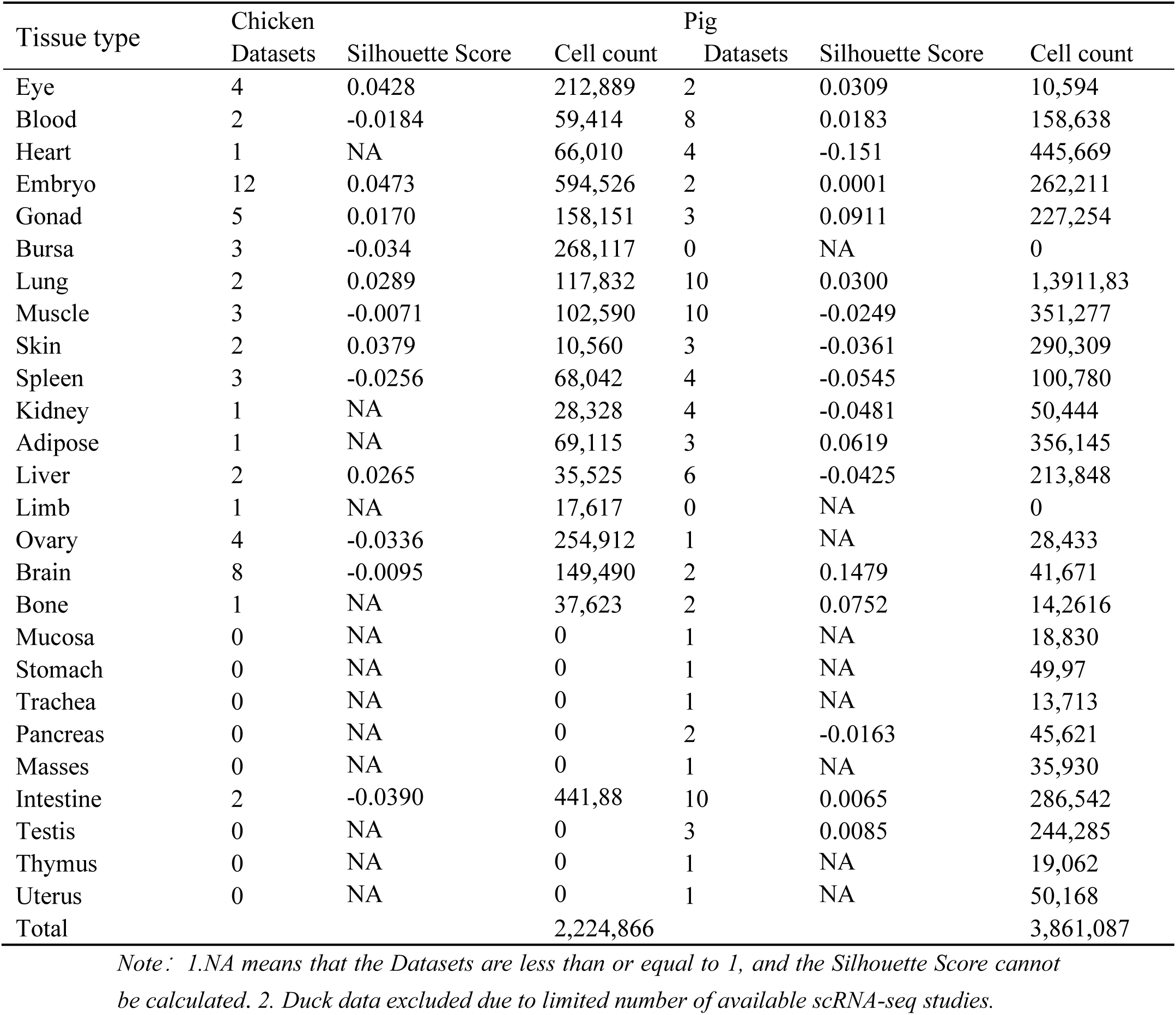
Data summary of Atlas Module.

In terms of functional expansion, AniMarkerDB supports multi-dimensional search, interactive visualization (including word clouds, bar charts, and heatmaps), as well as tools for cross-species marker gene comparison, protein structural similarity assessment, and user-defined cell annotation (Figure 1). The homepage summarizes coverage by species (Figure 2A) and visualizes usage trends via a Sankey plot (Figure 2B), together with panels that display the most-searched cell types and the top pathogens by epitope counts for each host species (Figure 2C). The resource is continuously updated to track new single-cell datasets and epitope records. The platform is continuously updated with newly published single-cell transcriptomic datasets and epitope information, ensuring alignment with the latest research developments.

**Figure 2.**
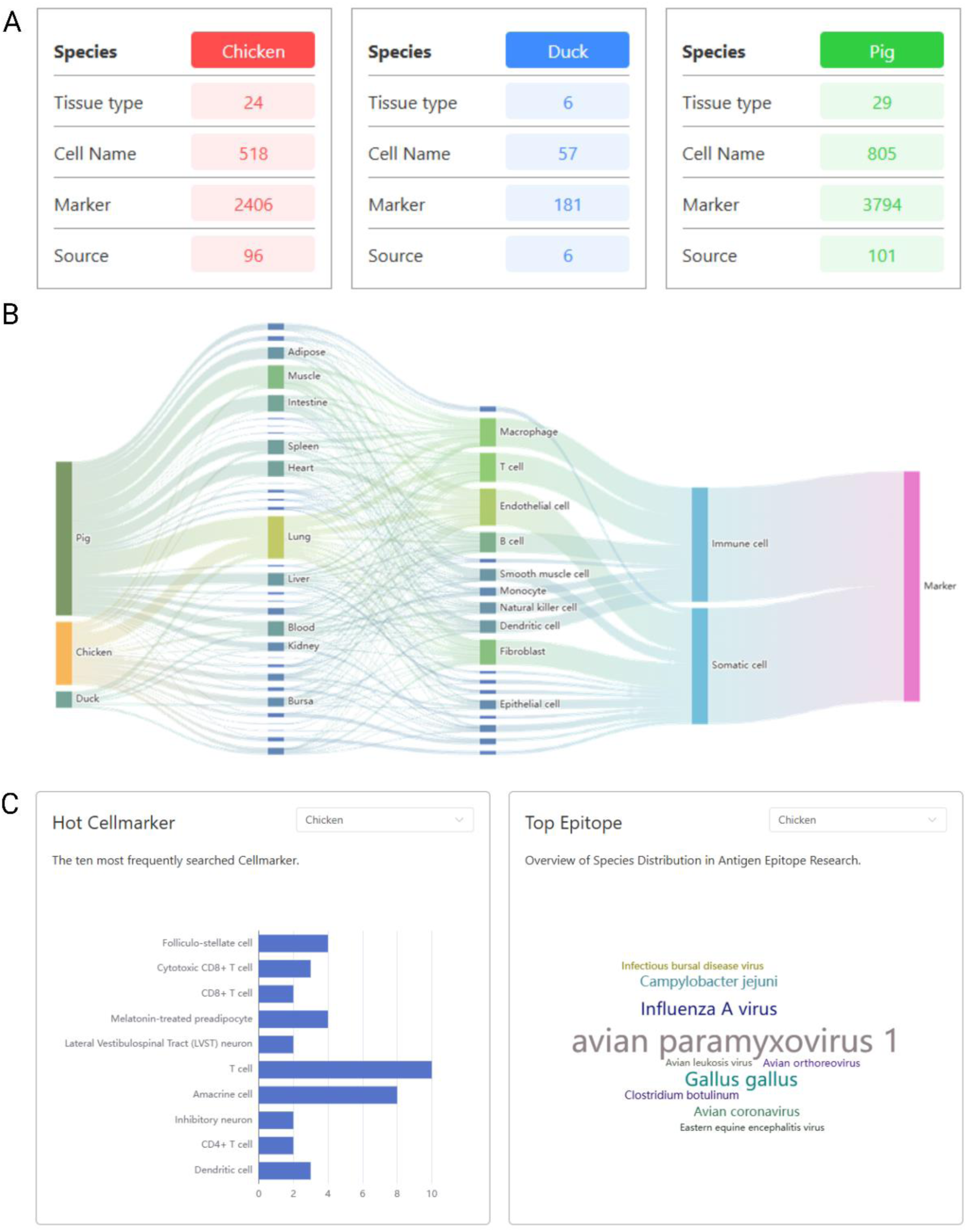
Distribution of marker and epitope data in AniMarkerDB. (**A**) Summary statistics of chicken, duck, and pig marker entries, including numbers of tissue types, cell types, markers, and literature sources. (**B**) Sankey diagram showing the relationships among species, tissues, cell types, and the immune/somatic classification of marker-associated cells. (**C**) Left: Top 10 most frequently searched cell types in the marker module; Right: The top 10 pathogens with the highest number of epitopes recorded for the selected host species.

AniMarkerDB makes extensive use of public resources by integrating not only epitope data from IEDB but also marker gene information for mouse and human from CellMarker 2.0. Epitope statistics showed that most entries were derived from pig, followed by chicken and duck, with peptide lengths predominantly distributed between 9 and 11 amino acids (Figure 3A), consistent with typical MHC-I studies. At the pathogen level, African swine fever virus (ASFV) and porcine reproductive and respiratory syndrome virus (PRRSV) accounted for the largest number of epitopes, while avian pathogens were mainly represented by avian paramyxovirus type 1 and influenza A virus (Figure 3B). Cross-species comparison of marker genes revealed that endothelial cells, macrophages, T cells, and fibroblasts exhibited the highest conservation (Figure 3C), with detailed shared markers illustrated in the heatmap (Figure 3D, showing genes present in ≥4 species). Additional statistics indicated that human and mouse shared the largest number of common cell types (656), followed by human and pig (224) (Figure 3E). Overall, these results demonstrate the broad coverage and systematic integration of epitopes and marker genes across species within the platform.

**Figure 3.**
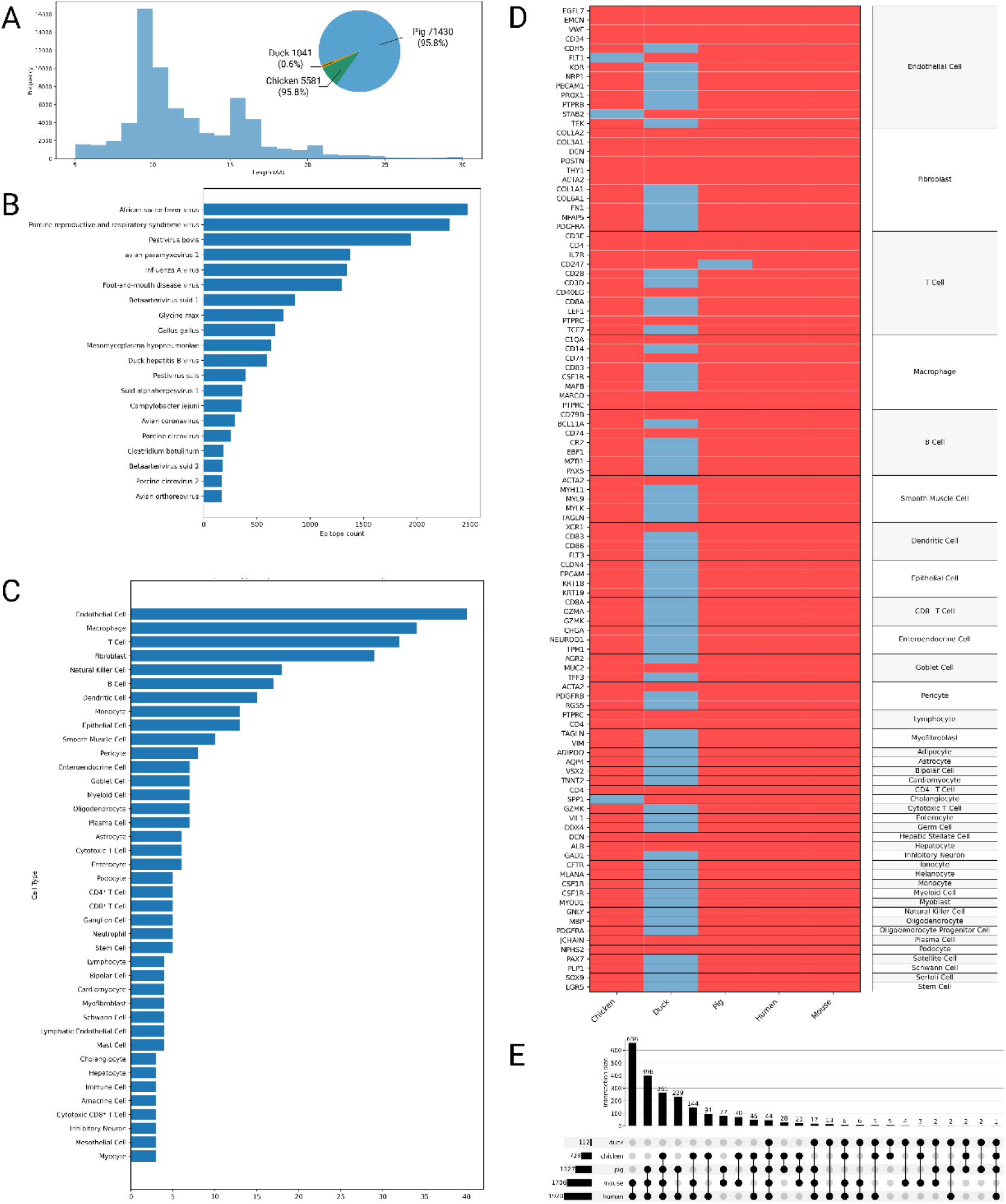
Epitope composition, pathogen distribution, and cross-species marker gene comparison in AniMarkerDB. (**A**) Histogram of epitope length distribution and pie chart of species composition. (**B**) Bar chart of epitope counts for different pathogens. (**C**) Bar chart of cell types ranked by the number of marker genes shared across three or more species. (**D**) Heatmap of cross-species marker genes across cell types. The heatmap shows conserved marker genes present in at least four species, with red indicating presence and blue (**E**) UpSet plot showing intersections of cell-type among species.indicating absence.

Taken together, the establishment of AniMarkerDB substantially enriches the landscape of systematic resources dedicated to single-cell marker genes and immune epitopes in poultry and livestock. Through rigorous standardization and a diverse array of functionalities, the platform enables high-resolution exploration of cellular heterogeneity, immune mechanisms, molecular breeding, and disease resistance in economically important animal species.

### User Interface and Application Scenarios of AniMarkerDB

AniMarkerDB features an intuitive and hierarchical web interface that supports rapid information retrieval and in-depth exploration of marker genes and immune epitopes for livestock and poultry research (Figure 4). Upon visiting the homepage, users can quickly search by entering keywords such as gene names, tissues, or cell types in the top search bar or by using the species selector. Below, an anatomical overview presents the major organ systems of species like chicken and pig as clickable nodes. Users can click different organ regions to quickly navigate to the summary page of marker genes for corresponding tissues and cell types, enabling direct visualization of the distribution of main tissue cells (Figure 4A). For example, when “chicken” is selected, clicking its icon displays a visual overview of key organs such as the bursa of Fabricius, thymus, and liver. By clicking on the relevant organ area, users can view statistics for all cell types within the tissue, the number of main marker genes, and a collection of related literature for that tissue (Figure 4B). In addition to anatomical navigation, AniMarkerDB offers robust custom cell annotation functionality directly on the homepage. In the “Cell Annotation” module, users can upload their own gene sets. The platform automatically performs enrichment analysis against all cell-type markers in the database. It outputs the top 10 most enriched cell types (displayed as bar or heat maps), while showing the marker overlap for each cell type. For example, when uploading T cell-associated genes such as CD3D and CD8A, the system highlights “CD8^+^ T cell” as the most probable match and simultaneously displays which input genes match the markers for this cell type (Figure 4C).

**Figure 4.**
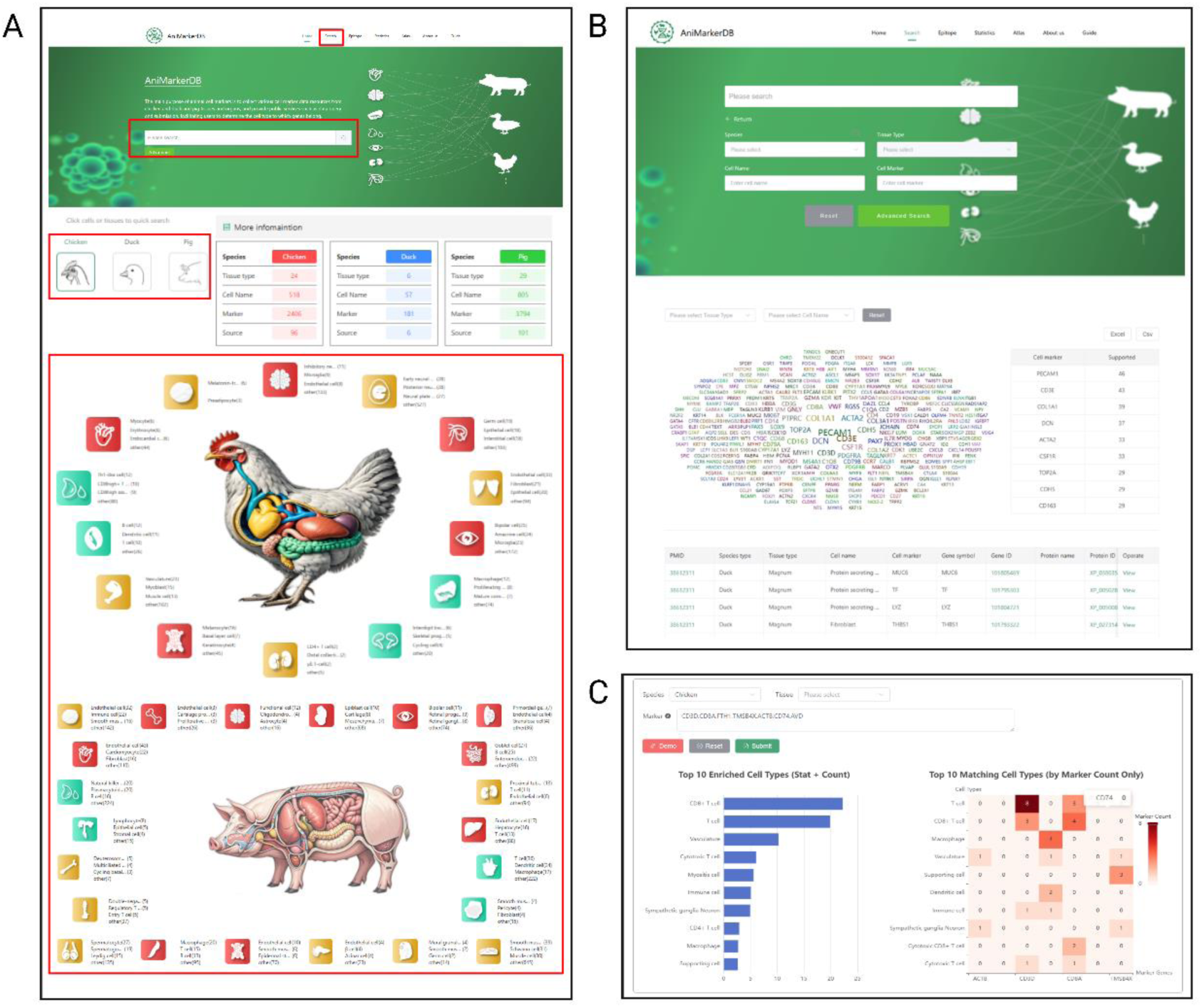
Overview of the AniMarkerDB interface and core functionalities. (**A**) The homepage interface integrates a quick search bar, species selector, and an anatomical overview of major tissues, each linked to corresponding cell marker entries, enabling quick access to the associated cell types. (**B**) The advanced search page allows queries by species, tissue, cell type, and gene, with results visualized as a marker word cloud and table. (**C**) The marker annotation module supports user-uploaded gene sets, returning predicted cell types ranked by enrichment scores and marker overlap, displayed via bar chart and matrix heatmap.

The “Advanced Search” page enables users to flexibly combine criteria such as species, tissue, cell type, and marker gene for refined queries. The search results are displayed as word clouds and tables, both sorted by the number of supporting publications. For instance, to systematically query “porcine lung T cells and their marker genes,” users can sequentially select “Species: Pig,” “Tissue: Lung,” and “Cell Type: T cell.” The platform will return word clouds and heatmaps of all T cell-related markers (Figure 5A). Users can further pinpoint specific cell types (*e.g*., “T cell”) in the list above the word cloud, with the platform then displaying the corresponding marker gene word cloud and ranking table (Figure 5B). By clicking any marker gene entry (such as CD3E), users are directed to the detailed gene information page, which includes the gene symbol, Gene ID, Protein ID, supporting PMID, and annotation of tissue/cell type, among other metadata (Figure 5C). Moreover, the immune epitope query function is a highlight of AniMarkerDB. Users can quickly switch to the “Epitope Module” from the homepage, which integrates IEDB immune epitope data. For immune cell-related marker genes, users can directly access the “Epitope Module” from the detailed gene information page to perform multidimensional interactive queries based on host species, pathogen, epitope type (MHC-I/II, etc.), and experimental category (Figure

**Figure 5.**
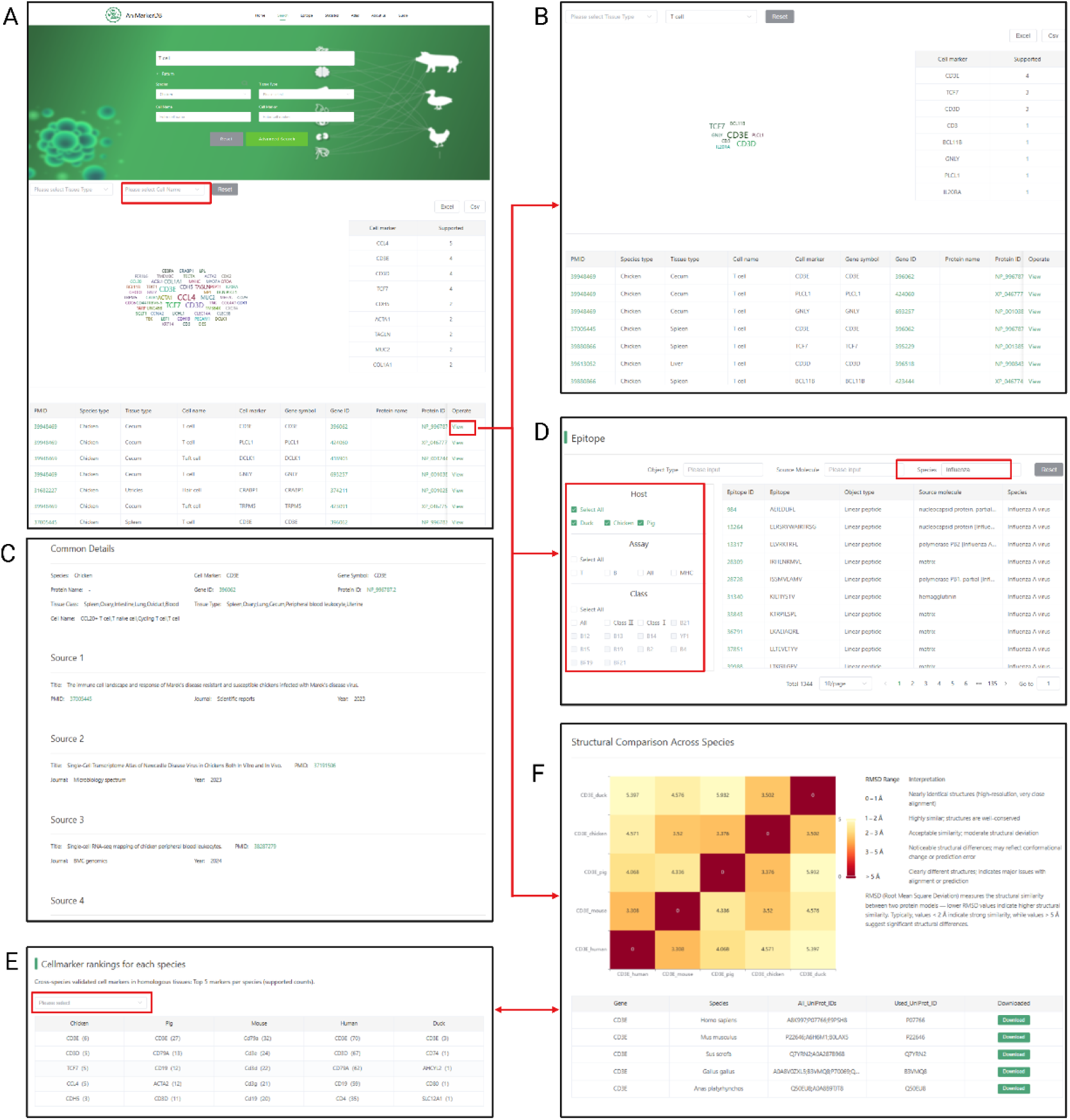
Search result pages of AniMarkerDB. (**A**) Marker gene search interface allowing queries by species, tissue, cell type, or gene symbol. Searches return multiple candidate cell types along with associated markers. (**B**) Refined search results for a selected cell type, highlighting relevant marker genes. (**C**) Detailed view of a specific marker gene, including gene symbol, protein IDs, and reference sources. **(D)** Epitope records associated with immune cell markers, with filtering options for host, pathogen, and assay type. (**E**) Cross-species comparison of the selected cell type, showing shared and species-specific marker genes among chicken, pig, duck, human, and mouse. (**F**) Structural comparison of marker gene proteins across species. A heatmap of RMSD values summarizes 3D protein similarity, with structure download links.

5D). All results support batch download, greatly facilitating applications such as infectious disease research and vaccine target design. The platform also supports systematic horizontal comparisons of cell types and their marker genes across species. The “Cross-species Comparison Module” can be found under the result panel of a particular cell type search; for example, selecting “T cell” will automatically list the top 5 markers in chicken, pig, duck, human, and mouse (the latter two derived from CellMarker 2.0), and identify common markers such as CD3D and CD3E across species (Figure 5E). For any given marker gene, users can directly access the “Cross-species Structure Comparison” on its detailed page and visualize 3D protein structure similarities via RMSD heatmaps (Figure 5F). For example, the details page of CD3E reveals that the greatest structural difference is between duck and human, with an RMSD value of 5.397. Such results provide evidence for studies of functional conservation and antibody cross-reactivity.

Additionally, the “Atlas Module” in AniMarkerDB enables the visualization of single-cell atlases and marker gene expression for specific tissues/cell types. Users can select target species, tissues, and cell types, and view the distribution of cell subpopulations in UMAP/t-SNE dimensionality reduction plots. For example, if researchers wish to examine the “spatial distribution and marker expression of CD4^+^ T cells in chicken lung,” they can simply select the target tissue and cell type in the Atlas interface (Figure 6A); the platform will automatically present the distribution of cell subgroups and the top expressed marker word clouds (Figure 6B and C), with the ability to hover for detailed metadata for each cell. Users can also click to examine the heatmap of a particular marker gene’s expression across cell types (Figure 6D). The Atlas module further integrates GSEA, GO, and KEGG enrichment analyses, as well as PPI networks (Figure 6 E, F and G), enabling users to analyze the function and regulatory networks of different cell types across various species and tissues in one step.

**Figure 6.**
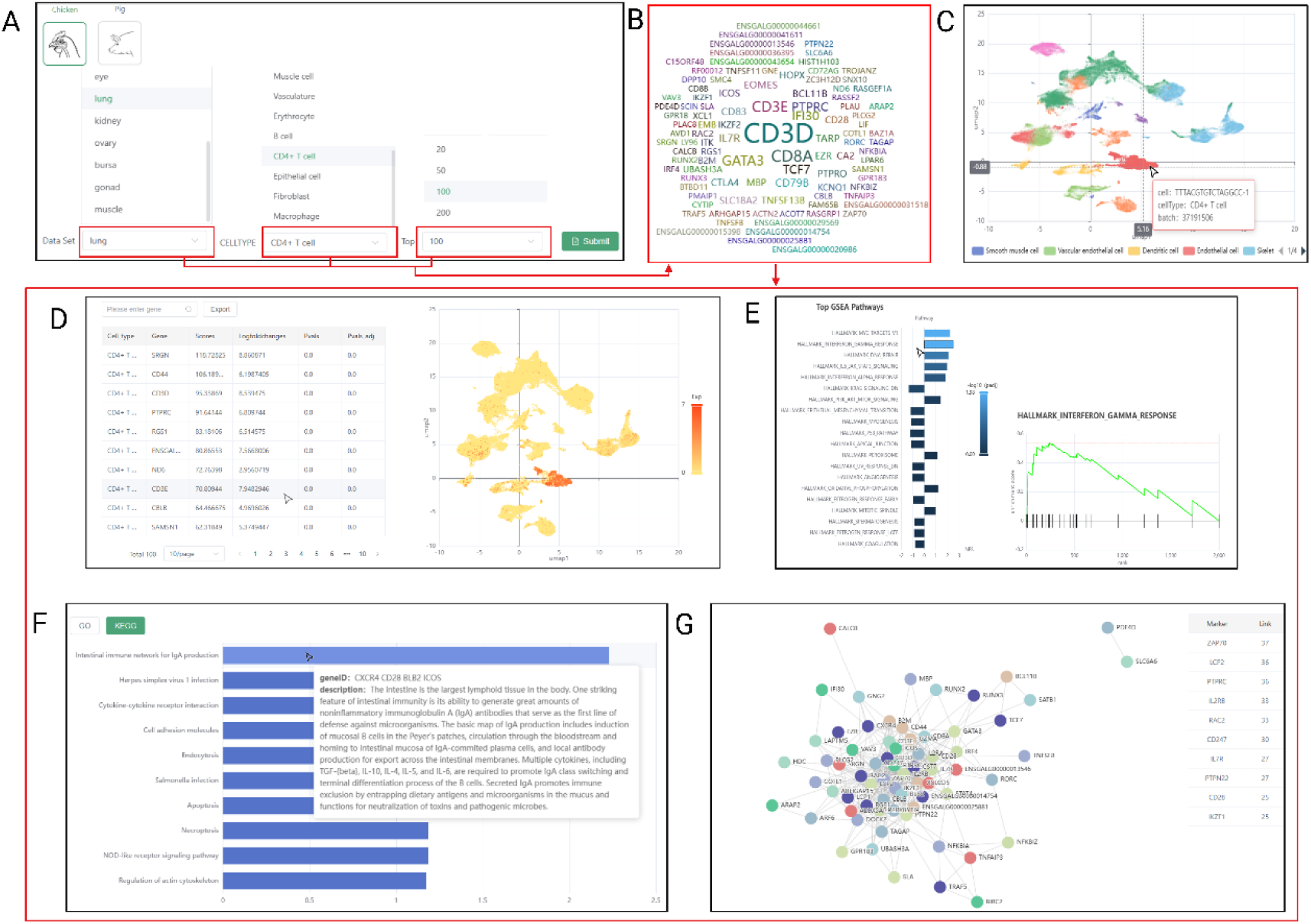
Functional characterization and visualization of cell types in tissue-level atlases. (**A**) Selection interface for selecting interest species, tissue, and annotated cell types for downstream analysis. (**B**) Word cloud visualization displaying the top marker genes for the selected cell type, which are subsequently used for downstream enrichment analyses. (**C**) UMAP visualization of the selected tissue, with cells colored by annotated cell types. Hovering reveals metadata for individual cells. (**D**) Ranked marker gene list for the selected cell type, alongside a feature plot showing the expression of a selected gene across all cells. (**E**) GSEA (Gene Set Enrichment Analysis) results for the selected cell type. (**F**) GO and KEGG enrichment results with toggle support; pathway descriptions and matching marker genes are shown interactively. (**G**) Protein– protein interaction (PPI) network constructed from top differentially expressed genes using STRING, with the 10 most connected proteins based on interaction degree.

### Database Management and Continuous Updates

AniMarkerDB is supported by a dedicated backend management system that enables real-time curation and maintenance of marker genes, immune epitopes, and single-cell atlases. The system provides secure administrator access(Figure 7A) and organizes all records in a standardized tabular format containing key metadata such as PMID, species, tissue type, cell type, sequencing technology, gene symbol, Gene ID, and Protein ID. Administrators can efficiently filter records, conduct batch imports or exports, and upload standardized datasets via pre-defined templates, ensuring consistency and accuracy. In addition to ad hoc updates(Figure 7B), AniMarkerDB follows a scheduled quarterly update cycle, during which the latest literature is systematically reviewed, newly released single-cell transcriptomic datasets are integrated, and relevant entries from external resources such as IEDB, CellMarker 2.0, and public repositories including GEO, GSA are incorporated. This continuous management and update mechanism ensures that AniMarkerDB remains comprehensive, current, and aligned with the latest advances in single-cell biology and immunological research in livestock and poultry.

**Figure 7.**
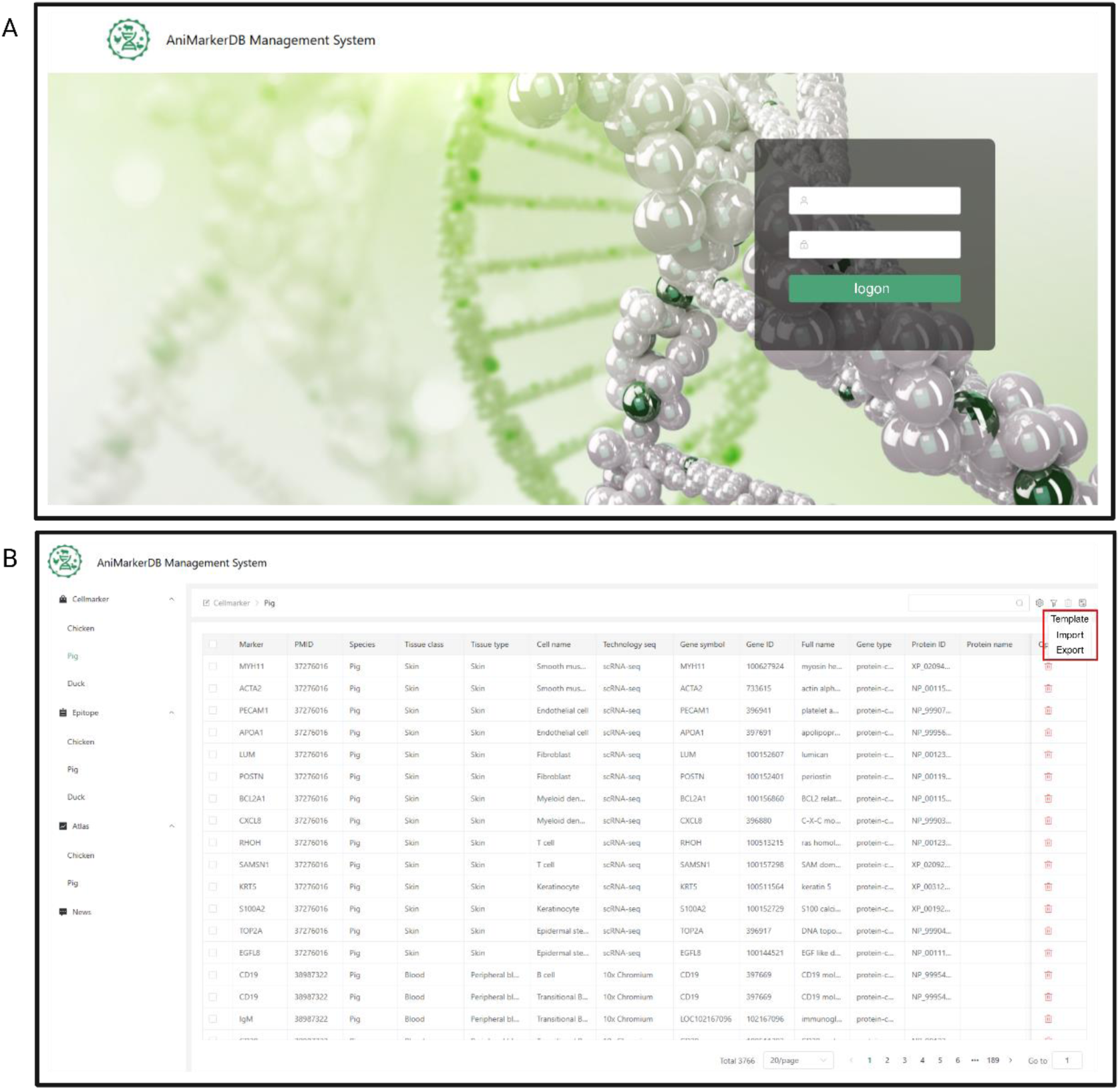
AniMarkerDB management system for database maintenance and updates. (A) Secure login interface for the AniMarkerDB backend management system. (B) Data management panel for browsing, editing, and batch import/export of marker genes, immune epitopes, and single-cell atlas records.

In summary, AniMarkerDB, through multi-level data retrieval, visualization, and functional expansion, enables users to explore from species and tissues down to cells, genes, and epitopes in a full pipeline manner. Whether for basic biological research, functional genomics, vaccine or antibody development, or rapid annotation and validation of single-cell data, AniMarkerDB provides researchers with an intuitive, systematic, and authoritative all-in-one resource and support.

## Discussion

The rapid proliferation of single-cell transcriptomic studies in livestock and poultry has produced a large volume of valuable data. However, the lack of standardized, high-coverage resources has significantly hindered downstream comparative and functional analyses (10,35). AniMarkerDB addresses this critical gap by systematically curating and integrating marker genes and immune epitopes from economically significant species—such as chicken, pig, and duck— into a unified, queryable framework.

First, AniMarkerDB currently provides the most comprehensive marker gene resource for livestock and poultry, systematically integrating 6352 high-quality marker gene records across more than 26 tissues and 791 cell types. This extensive data foundation enables robust horizontal and vertical analyses across multiple tissues, cell types, and species, significantly broadening the scope and depth of animal cell atlas research. These features allow researchers to perform reliable comparative studies while minimizing issues caused by inconsistent nomenclature or fragmented annotations—challenges that are still prevalent in livestock scRNA-seq literature. Beyond its comprehensive data coverage, a key strength of AniMarkerDB lies in its multi-layered standardization pipeline, which includes harmonized gene symbols, cell ontology alignment, and rigorous metadata curation. This ensures consistency and interoperability across datasets, enabling users to conduct reliable comparative studies without being hindered by inconsistent annotations or fragmented records—issues that remain prevalent in livestock scRNA-seq literature. (10,11). Furthermore, integrative analyses of epitope composition and marker gene conservation underscore the database’s ability to capture both species-specific molecular features and cross-species commonalities, thereby validating its utility for comparative immunogenomics.

In addition, AniMarkerDB offers a flexible and powerful suite of analytical tools designed to support the diverse research needs of the livestock and poultry research communities.. These tools include cross-species marker gene comparison, protein structural similarity analysis, immune epitope retrieval, visualization of single-cell atlases, functional enrichment, and user-defined annotation. Users can easily retrieve marker genes specific tissues or cell types, perform multi-species functional and evolutionary analyses, and support applications in basic research, molecular breeding, and disease prevention. These capabilities not only facilitate basic and translational research, but also enhance molecular breeding and disease prevention strategies, particularly in non-model species. Moreover, the platform supports interactive data visualization and export, reducing technical barriers and promoting reproducibility in single-cell workflows.

Despite its current strengths, AniMarkerDB remains limited in species coverage, focusing primarily on chicken, pig, and duck. However, the critical importance of these economically valuable animals in agriculture, food security, and as models for zoonotic disease research underscores the urgent need for systematic resources in this field(36). With the rapid advancement of single-cell sequencing and spatial multi-omics technologies, the capacity to unravel cellular complexity and disease mechanisms in non-model and emerging animal species is expanding, making comprehensive marker gene databases increasingly indispensable(37). To ensure sustainability and data freshness, AniMarkerDB is supported by a linked management system that coordinates literature tracking, data ingestion, quality control, and version updates. This system not only facilitates timely incorporation of new single-cell and epitope datasets but also provides a framework for user contributions under expert curation, thereby maintaining long-term reliability and community engagement. Accordingly, future development of AniMarkerDB will focus on broadening its species scope to encompass additional livestock, poultry, and wild animals of agricultural and biomedical significance, while enriching the diversity and resolution of data modalities and annotations. In particular, integrating single-cell datasets from both healthy and pathogen-infected animals will provide valuable resources for the study of animal disease models and host-pathogen interactions, ultimately contributing to the prevention and control of zoonotic diseases. The platform’s adherence to FAIR principles and collaboration with international consortia will further enhance interoperability, sustainability, and global impact. Taken together, AniMarkerDB is positioned as a key resource to support single-cell research in non-model and economically important animals, and will continue to play a pivotal role in advancing functional genomics, disease resistance, and translational research in agricultural and veterinary sciences.

In conclusion, AniMarkerDB represents a foundational advance in livestock and poultry single-cell research. By integrating high-confidence molecular signatures and immune features into a unified, user-friendly platform, it enables the scientific community to explore cellular complexity, functional divergence, and immunological mechanisms with unprecedented resolution. We anticipate that AniMarkerDB will serve as a pivotal enabler for both fundamental discovery and applied innovations in animal health, breeding, and disease resilience.

## DATA AVAILABILITY

AniMarkerDB can be accessed at https://animarkerdb.bio

## AUTHOR CONTRIBUTIONS

Zhuohang Li: Writing—original draft, Formal analysis, Data curation, Conceptualization. Tao Zhang: Data curation, Formal analysis. Xueqing Li: Writing—review & editing. Jiangwu Huang: Data curation. Zimin Xie: Data curation. Fei Gao: Data curation. Haiming Cai: Funding acquisition. Mingfei Sun: Funding acquisition. Manman Dai: Conceptualization, Writing—review & editing, Funding acquisition. Ming Liao: Funding acquisition.

## ACKNOWLEDGEMENTS

We acknowledge the performance computing resources support from Guangzhou Minglead Gene Technology Co., Ltd. in the development and maintenance of the platform. We are grateful to all the data contributors whose invaluable contributions have made this project possible.

